# Using integrated wildlife monitoring for quantifying the impact of emerging Bagaza virus in red-legged partridges in Portugal, 2012-2024

**DOI:** 10.1101/2024.08.07.606962

**Authors:** João Queirós, João Basso Costa, Gonçalo de Mello, Catarina Fontoura-Goncalves, Úrsula Höfle, Christian Górtazar, Paulo Célio Alves, Pelayo Acevedo, David Goncalves

## Abstract

Bagaza virus, a vector-borne flavivirus that causes significant mortality in wild bird species, emerged in Portugal in September 2021. This study used integrated wildlife monitoring to quantify its impact on a natural population of red-legged partridge (*Alectoris rufa*) in southern Portugal. We constructed a baseline model of population dynamics prior to the outbreak (2012-2021) and used this model to predict population size in subsequent years (2022-2024). This model, which included population abundance and demographic data of red-legged partridge, as well as climatic variables, showed a strong fit to the observed data, and the difference between the predicted and observed population size in 2022 suggests a Bagaza-induced mortality rate of at least 59% (95% CI 52-66%). In addition to enabling the adaptive management of populations, our results demonstrate that integrated wildlife monitoring offers epidemiological insights for risk assessment, highlighting its role in proactive wildlife disease surveillance within the One Health framework.

## Introduction

Emerging pathogens pose significant global health concerns, causing substantial morbidity and mortality impacts in humans and animals (1). While identifying and quantifying disease-induced mortality is relatively straightforward in humans and domestic animals, assessing their effects on wildlife remains challenging (2) due to the inherent difficulties in monitoring population dynamics and health status in natural settings (3). To bridge this gap, integrated wildlife monitoring (IWM) has emerged as a crucial tool within the One Health approach, enabling the timely detection of emerging pathogens in natural populations and tracking changes in disease dynamics across diverse host-pathogen-environment contexts by quantifying host population characteristics (4).

By monitoring host abundance, demography and health simultaneously, IWM facilitates early threat identification and provides valuable insights into the effects of emerging pathogens on wildlife populations (5). These insights are vital for assessing potential spillover risks to domestic animals or human populations. However, effective wildlife population monitoring requires sustained long-term efforts and standardized methodologies (6).

Bagaza virus (BAGV) is an emerging vector-borne arbovirus in Europe and other parts of the world (7). It emerged in Portugal in September 2021, associated with an outbreak of the disease in red-legged partridge (*Alectoris rufa*), but has also affected other wild birds (8). BAGV is a positive-sense single-stranded RNA virus belonging to the mosquito-borne cluster of the genus flavivirus, which includes other pathogenic viruses such as dengue, West Nile, Zika, Japanese encephalitis, and yellow fever viruses, all associated with neurological disease in wild and domestic animals as well as humans (9). Based on enquiry surveys, a retrospective study of the first BAGV outbreak in Spain in 2010 estimated a mortality rate of 23% in red-legged partridge populations (10), while a controlled experimental study estimated a mortality rate of 30% (11). However, empirical data to quantify the impact of this mosquito-borne flavivirus on natural red-legged partridge populations is currently lacking. The red-legged partridge is a game species of significant ecological, economic and social importance in southern Europe. While more than four million birds are reared in intensive farms and released annually in hunting grounds in Portugal and Spain (12), natural populations have been declining in recent decades, threatening the sustainable management of this natural resource and, ultimately, the conservation of this species in the wild (13).

This study aims to quantify the impact of the 2021 BAGV outbreak on the dynamics of a red-legged partridge population in southern Portugal. Using empirical data on host population abundance, demography, and climatic variables collected between 2012 and 2024, we constructed a baseline model of population dynamics prior to the outbreak (2012-2021) and used this model to predict population size in subsequent years (2022-2024), particularly in January 2022, immediately after the BAGV circulation in this population. We showed that IWM can effectively estimate the impact of an emerging pathogen within a wild population.

## Material and methods

### Study site, demographic monitoring and climatic variables

The study was conducted in southern Portugal at “Herdade de Vale de Perditos” in Serpa (37.82236N, -7.37953W). This 3,000-hectare hunting ground features a Mediterranean climate and vegetation. The terrain is sloping, and active landscape management is employed to enhance habitat suitability for the red-legged partridge. The estate’s mosaic landscape includes cultivated parcels intermingled with natural shrubs and cork (*Quercus suber*) and holm oak (*Quercus ilex*) trees. Additionally, more than 200 artificial feeding and watering stations are provided year-round throughout the estate to supplement the partridge’s diet and water resources, which are particularly important during the summer. Since 2012, due the implementation of monitoring program, regular population size estimations of the red-legged partridge have been conducted three times every year: i) in January, after the hunting season (Jan_pop_size); ii) in March, during the breeding season and when most of the birds are mated (Mar_pop_size); and iii) in July (Jul_pop_size), after the breeding season. In January and March, the partridge abundance is estimated by dividing the area into nine units and counting individuals by walking across these units – a line of six beaters walks the sampled unit and registers the partridges observed, a sort of blank beat/strip census (14,15).

In July, in each of the nine units, adults and juveniles (mostly 12 < weeks > 8) are counted along transects (about 15 km each) made by car (in both directions, on different days) to calculate the age ratio (juveniles/adults) (16,17). Age ratio is further used to estimate the number of juveniles in July, using July’s adult estimation as a reference. Adult numbers in July are calculated using March population size estimations and assuming a monthly mortality rate associated with predation of 5% (from March to July) (18). We considered the minimum mortality rate reported in adult red-legged partridge in Iberia (18) as intense predation control is regularly performed in this hunting ground. The population size in July is inferred by summing estimated adults and juveniles. Climatic variables, including the monthly average of the daily mean temperature and cumulative monthly rainfall, are primarily compiled from an in-house meteorological station supplemented with data from the nearest national meteorological station, that of Beja. Consistent application of standardized methodology and management practices over the years reduces potential uncertainties associated with the counting estimates of population size.

### Population dynamics model

We developed a baseline Generalized Linear Model (GLM) using negative binomial regression with a logit link to model the dynamics of the red-legged partridge population. This model used ten years of demographic data collected before the BAGV outbreak, which occurred between September and November 2021 (8). The aim was to predict the population size in January for the subsequent years of 2022, 2023, and 2024. Discrepancies between the predicted and observed population sizes in 2022 can be primarily attributed to the impact of the BAGV outbreak, as management practices remained consistent. However, for 2023 and 2024, no deviations were anticipated, as BAGV appeared to no longer be circulating within the population (Fontoura-Gonçalves et al, submitted). The GLM was designed to adjust the population size in January, after the hunting season (October - December) and before the reproductive season (February - June), based on a simple population dynamics model of Birth-Immigration-Death-Emigration (BIDE) (19). However, dispersal (emigration/immigration) and productivity (nest success/chick survival) in Mediterranean ecosystems are deeply influenced by the winter/spring climate, which modulates the habitat conditions required for nesting and the diet of chicks, especially during the first month of life (18). As these parameters are difficult to assess accurately in natural populations and require complex integrated population modelling approaches (20), our population dynamics model avoids using them by considering the population size estimated in July (Jul_pop_size). This parameter already considers dispersal events, as it is derived from the population size in March, after the birds have formed pairs and established a territory, and productivity effects in the population, as most birds with 8 to 12 weeks of age have already passed the peak of mortality (18). Between July and December, we assumed no emigration/immigration and considered a monthly mortality primarily induced by predation (Nat_mortality) of 5% for adults and 10% for juveniles (sub-adults), as survival of sub-adults is usually low (18). Mortality induced by hunting (Hunting_bag) is recorded during each hunting season (October-December). To reduce collinearity between these variables (Jul_pop_size, Nat_mortality, Hunting_bag), assessed through the variant inflation factor (*vif*) function from the package *car* (21), we conducted a principal component analysis (PCA) using the prcomp function from the multcomp package (22). The resulting three principal components (PC1, PC2, and PC3) explained all observed variance (Table S1) and contributed to improving the model fit. Additionally, we explored the influence of the climatic variables monthly average of the daily mean temperature and cumulative monthly rainfall recorded between July and December. The GLM was parameterized using the glm.nb function from the MASS package (23), and model predictions along with 95% confidence intervals were generated using the add_ci function from the ciTools package (https://github.com/jthaman/ciTools). Visualization of the variation along years of the observed, fitted and predicted population sizes from the GLM was done using the ggplot function from the ggplot2 package (24). All analyses were conducted within the R environment (25).

## Results

Predictions beyond the baseline model from 2022 to 2024 yielded divergent outcomes. In 2022, following the 2021 BAGV outbreak, the predicted population size (January abundance) was 6380 individuals (95% CI 6105 – 6667), which exceeded the observed size by 66% (95% CI 59.1 – 73.7%) (Figure 1).

**Figure 1:**
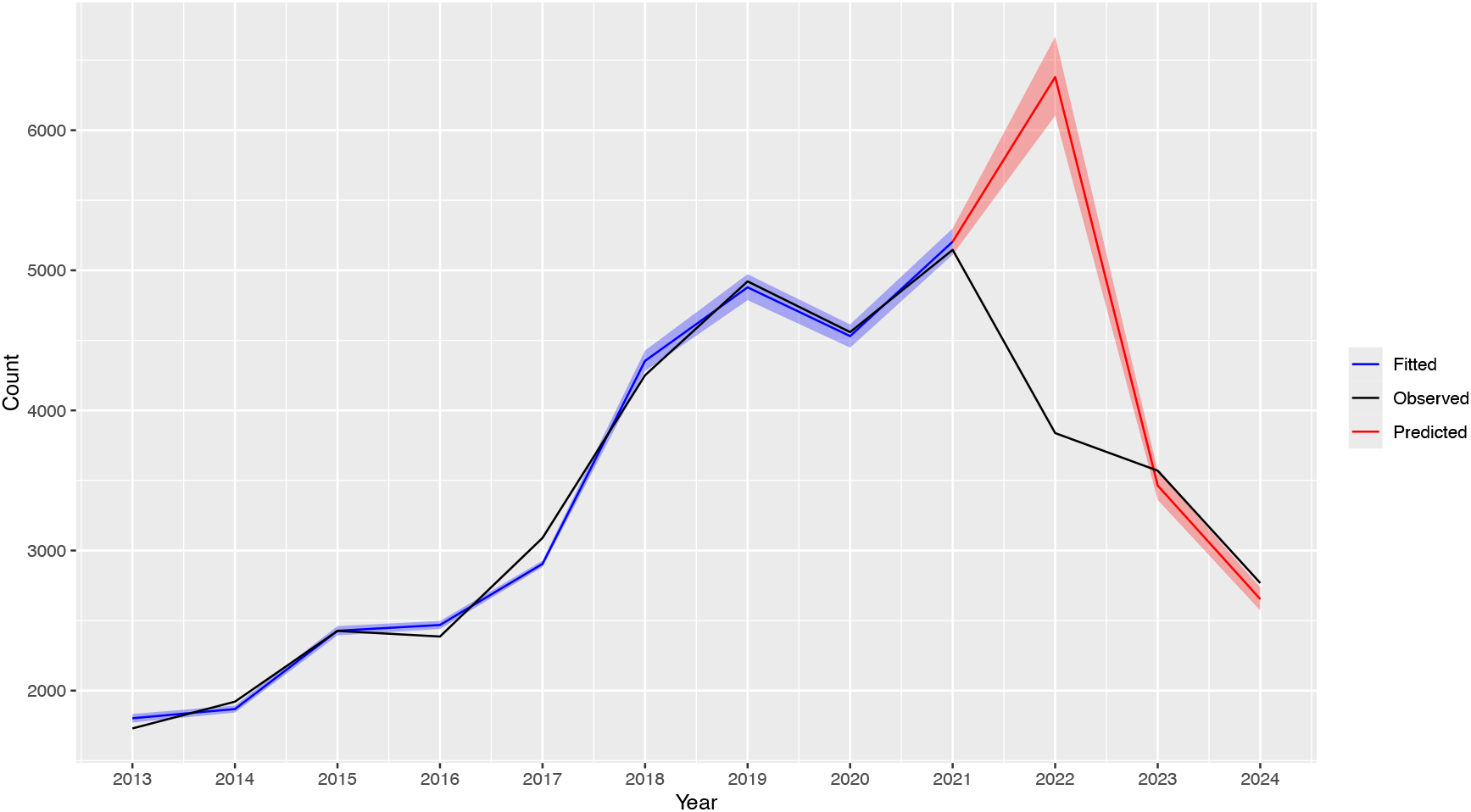
Red-legged partridge population size observed (black line), fitted (blue line) and predicted (red line) in January between 2013 and 2024. Predictions were based on a Generalized Linear Model that used 10 years of demographic data (2012-2021) collected before the 2021 BAGV outbreak. Blue and red shallows represent the 95% confidence intervals for the fitted and predicted values, respectively.

The corresponding predicted density in 3,000 hectares was 2.13 individuals per hectare (95% CI 2.03-2.22%), which would be the highest value in January for the study period but still lower than the densities recorded in July between 2017 and 2021 (Table 1). Conversely, in 2023 and 2024, the predicted values fell below the observed values by 3.0% (95% CI 0.1 – 5.8%) and 4.2% (95% CI 1.1 – 7.1%), respectively (Figure 1). After subtracting the maximum standard error of ∼7% (95% CI) observed in the model prediction, we estimated a BAGV-induced mortality rate of at least 59% (95% CI 52 – 66%).

**Table 1:**
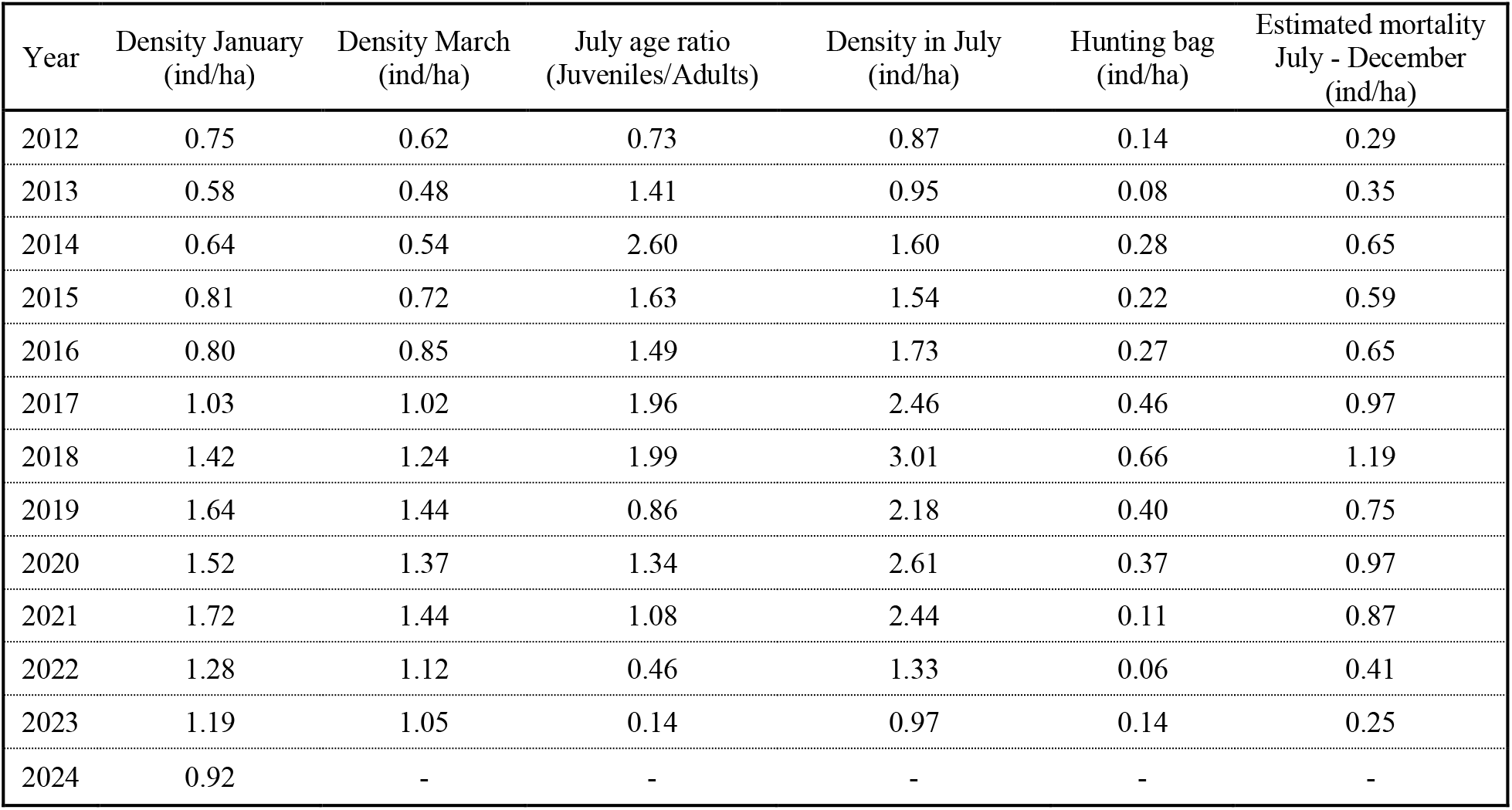
Demographic parameters estimated (observed) for the red-legged partridge population between 2012 and 2024.

| Year | Density January<br>(ind/ha) | Density March<br>(ind/ha) | July age ratio<br>(Juveniles/Adults) | Density in July<br>(ind/ha) | Hunting bag<br>(ind/ha) | Estimated mortality<br>July - December<br>(ind/ha) |
| --- | --- | --- | --- | --- | --- | --- |
| 2012 | 0.75 | 0.62 | 0.73 | 0.87 | 0.14 | 0.29 |
| 2013 | 0.58 | 0.48 | 1.41 | 0.95 | 0.08 | 0.35 |
| 2014 | 0.64 | 0.54 | 2.60 | 1.60 | 0.28 | 0.65 |
| 2015 | 0.81 | 0.72 | 1.63 | 1.54 | 0.22 | 0.59 |
| 2016 | 0.80 | 0.85 | 1.49 | 1.73 | 0.27 | 0.65 |
| 2017 | 1.03 | 1.02 | 1.96 | 2.46 | 0.46 | 0.97 |
| 2018 | 1.42 | 1.24 | 1.99 | 3.01 | 0.66 | 1.19 |
| 2019 | 1.64 | 1.44 | 0.86 | 2.18 | 0.40 | 0.75 |
| 2020 | 1.52 | 1.37 | 1.34 | 2.61 | 0.37 | 0.97 |
| 2021 | 1.72 | 1.44 | 1.08 | 2.44 | 0.11 | 0.87 |
| 2022 | 1.28 | 1.12 | 0.46 | 1.33 | 0.06 | 0.41 |
| 2023 | 1.19 | 1.05 | 0.14 | 0.97 | 0.14 | 0.25 |
| 2024 | 0.92 | - | - | - | - | - |

The estimated baseline demographic model (2012-2021) showed a strong fit to the data (McFadden’s R-squared = 0.994), incorporating the principal components representing the July population size, predation and hunting-induced mortality (July-December), and October’s average of the daily mean temperature (Temp_oct) (Table S1). These variables exhibited minimal collinearity (VIF: 1.1-1.7, Supplementary Table S2) and all significantly improved the model fit (Table 2). Minor discrepancies between fitted and observed population sizes occurred, peaking in 2017 (6.0%, 95% CI 5.2-6.7%, Figure 1).

**Table 2:** Results of the Generalized Linear Model (GLM, negative binomial with logit link) performed to model the red-legged partridge population size in January using 10 years of demographic data (2012-2021) collected before the BAGV outbreak. The principal components (PC1, PC2 and PC3) represent the estimated population size in July, the estimated natural mortality between July and December and the hunting bag (October-December). Temp_oct represents the average of the daily mean temperature of October.

| Variables | Estimate | Standard Error | z-value | *p-value* |
| --- | --- | --- | --- | --- |
| Intercept | 8.10 | 0.013 | 610.2 | <0.001 |
| PC1 | -0.19 | 0.007 | -29.0 | <0.001 |
| PC2 | -0.32 | 0.037 | -8.9 | <0.001 |
| PC3 | 1.49 | 0.117 | 12.7 | <0.001 |
| Temp_oct | 0.05 | 0.011 | 4.3 | <0.001 |

Over the study period, population density (individuals per hectare) in January ranged from 0.58 in 2013 to 1.72 in 2021, while in March, it varied from 0.48 in 2013 to 1.44 in 2019 and 2021 (Table 1). In July, the age ratio (juveniles/adults) varied between 2.60 in 2014 to 0.14 in 2023, and the population density ranged from 0.87 in 2012 to 3.01 in 2018 (Table 1).

## Discussion

Our findings represent the first empirical study documenting the impact of BAGV on a natural population of red-legged partridges. Even allowing some bias associated with model fitting (∼7% maximum), we estimated a BAGV-induced mortality rate of at least 59% (95% CI 52 – 66%), surpassing previous estimates for natural populations in southern Spain (23%) obtained from enquiry surveys (10) by more than twofold. It also exceeded the 30% mortality observed in the experimental infection of red-legged partridges with BAGV (11), suggesting that birds are more susceptible to morbidity effects in natural environments than in controlled conditions.

In July 2021, the population estimate was 2.44 individuals per hectare, which, although indicating a high population density (25), was lower than the values reached in 2017 (2.46), 2018 (3.01) and 2020 (2.61; Table 1). As no BAGV circulation was detected in this population before September 2021 and after December 2021 (26), there does not seem to be a direct relationship between population density in July and virus emergence in autumn, probably due to the complex interactions between environmental and climatic conditions, vectors and host populations. However, a density-dependent effect on the transmission dynamics of this virus is expected, as direct transmission between red-legged partridges has been demonstrated under experimental conditions (11). Additional data on vector community dynamics, climate and virus phylogeography would help to understand the reason for its emergence in September 2021 and further circulation until November 2021 (26). In addition, although October’s average of the daily mean temperature (Temp_oct) enhanced our baseline model fit, the reasons for this positive effect on host population size are unknown and need to be investigated.

The sudden population decline caused by BAGV has resulted in significant economic and social repercussions on the species conservation and hunting sector, given the species’ status as one of the most emblematic and sought-after small game birds in the Iberian Peninsula (12). In the studied population, this decline was mitigated by a reduction in hunting efforts of over 70% compared to the average yield in the four seasons preceding the outbreak (2017-2020, Table 1). Such informed baseline decisions are only achievable through IWM and were instrumental for the sustainability and conservation of this natural population. Subsequent years (2022 and 2023) witnessed minimal recruitment of juveniles into the population, with an age ratio (juvenile/adult) falling below 0.5 in July, hindering population recovery. However, similar management measures were not universally applied across the species’ range, jeopardizing the conservation of a species already experiencing a declining trend in natural populations in recent decades (13).

In addition to guiding wildlife management and conservation efforts, IWM provides invaluable epidemiological data for risk assessment analyses, particularly for pathogens with zoonotic potential like BAGV (4). Continued monitoring of this emerging pathogen in natural populations and at the interface with domestic animals and humans is imperative to understand its dynamics, including reasons for its emergence and disappearance, host range, and temporal and spatial impacts on host populations (5, 6).

In conclusion, our study emphasizes the critical role of IWM in elucidating the dynamics of emerging pathogens, thereby enhancing preparedness for potential interspecies transmission events to domestic animals or humans. It also reinforces the pivotal role of IWM within the One Health framework, offering a proactive approach toward averting future pandemics.

## Acknowledgements

We are grateful for the support of Herdade de Vale de Perditos and its staff for their help in collecting the data.

