## Appendix information for "Using integrated wildlife monitoring for quantifying the impact of emerging Bagaza virus in red-legged partridges in Portugal, 2012-2024"

**Table S1**: Factor loading of each demographic variable to the three principal components (PC1, PC2 and PC3) calculated in the Principal Component Analysis (PCA) and respective variance explained. The demographic variables included the estimated population size in July (Jul_pop_size), the estimated natural mortality between July and December (Nat_mortality) and hunting yield during the hunting season (Hunting_bag).

| Variables used in the PCA | PC1 | PC2 | PC3 |
|---|---|---|---|
| Jul_pop_size | -0.592 | -0.413 | 0.692 |
| Nat_mortality | -0.597 | -0.352 | -0.721 |
| Hunting_bag | -0.541 | 0.840 | 0.038 |
| Proportion of variance | 0.902 | 0.092 | 0.006 |
| Cumulative proportion of variance | 0.902 | 0.994 | 1.000 |

**Table S2**: Variant Inflate Factor (VIF) values observed for the explanatory variables included in the Generalized Linear Model (GLM).

| | Explanatory variables included in GLM | | | |
|---|---|---|---|---|
| | PC1 | PC2 | PC3 | Temp_oct |
| VIF | 1.29 | 1.05 | 1.71 | 1.46 |
